# Solvation Free Energy in Governing Equations for DNA Hybridization, Protein–Ligand Binding, and Protein Folding

**DOI:** 10.1101/2024.03.15.585270

**Authors:** Caroline Harmon, Austin Bui, Jasmin M. Espejo, Marc Gancayco, Jennifer M. Le, Juan Rangel, Daryl K. Eggers

## Abstract

This work examines the thermodynamics of model biomolecular interactions using a governing equation that accounts for the participation of bulk water in the balanced reaction. In the first example, the binding affinities of two model DNA duplexes, one of nine and one of ten base pairs in length, are measured and characterized by isothermal titration calorimetry as a function of concentration. The results indicate that the change in solvation free energy that accompanies duplex formation (Δ*G*^S^) is large and unfavorable. When normalized to the number of base pairs, the duplex with the larger number of G:C pairings yields the largest change in solvation free energy, Δ*G*^S^ = +460 kcal/mol/base pair at 25 °C. A modelling study demonstrates how the solvation free energy alters the output of a typical titration experiment. Hybridization measurements are completed at four different temperatures, however a van’t Hoff analysis of the data is complicated by the varying degree of intramolecular base stacking within each DNA strand as a function of temperature. The same thermodynamic framework is applied to a model protein– ligand interaction, the binding of ribonuclease A with the nucleotide inhibitor 3’-UMP, and to a conformational equilibrium, the change in tertiary structure of α-lactalbumin in molar guanidinium chloride solutions. The ribonuclease study yields a value of Δ*G*^S^ = +160 kcal/mol, whereas the folding equilibrium yields Δ*G*^S^ ≈ 0, an apparent characteristic of hydrophobic interactions. These examples complement previous applications of the governing equation to the interaction of smaller molecules and demonstrate the importance of solvation energy in biothermodynamics.

## INTRODUCTION

The stable binding of two molecules to form an active complex is a fundamental step in many biochemical reactions. Unfortunately, the classical thermodynamic equations for binding equilibria in water may not account fully for the contribution of the solvent. Because a complex between two molecules will interact with fewer water molecules than the unbound reactants, and because the chemical potential of a water molecule depends on the chemistry of any neighboring reactants or secondary solutes, changes in the free energy of water are expected to play a significant role in the thermodynamics of binding and conformational equilibria.

In 2013, Castellano and Eggers introduced a modified equation for binding equilibria that was derived from the chemical potential of all reaction participants, including the solvent.^1^ The thermodynamic framework, elaborated further in a subsequent paper,^2^ captures the change in free energy associated with the subset of water molecules that are released from the surfaces of the reactants upon binding. The framework has been applied to multiple host–guest pairings,^2,3^ in addition to evaluating the effects of secondary solutes on binding equilibria.^4^

The thermodynamic framework is summarized by Equations 1-5 below. For a binding reaction between two molecules, A and B, the following relationship is obtained for the overall change in reaction free energy when water is considered explicitly:

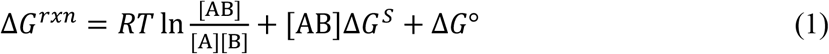

where R is the gas constant, T is the temperature in Kelvin, [AB] is the concentration of binary complex, Δ*G*^S^ is the change in solvation free energy, and Δ*G*° is the standard-state free energy constant. In general, the value of Δ*G*^S^ reflects all linked reaction equilibria that are not represented by the concentration ratio of products to reactants.^4^ For the current study, Δ*G*^S^ is expected to be dominated by the solvent-associated term, denoted Δ*G*^H2O^ in some of our previous work, and defined by the following:

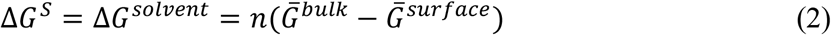

where “*n*” is the number of water molecules released per mole of complex formed, and where the bar above 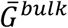 and 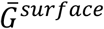 is a reminder that these represent location-averaged free energy values for a single water molecule in the bulk phase or next to a reactant surface, respectively. Because “*n*” is defined as the number of water molecules released per mole of formed complex, AB, the total contribution of water to the reaction free energy change is given by the term [AB]·Δ*G*^S^, as found in Equation 1. Of special note, the presence of 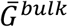 in the definition of Δ*G*^S^ (Equation 2) enables one to account for changes in equilibria due to the addition of secondary solutes that alter 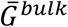 through cosolute–water interactions.^4^

The standard-state free energy in Equation 1, Δ*G*°, is obtained from combining all of the constant terms in the derivation that arise from the traditional reactants, leading to the following:

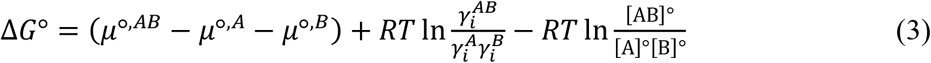

where *μ*°^,x^ are the standard-state potentials, 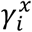 are the activity coefficients in solution *i*, and [x]° are the reference concentrations of each species *x*. Note that 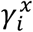 and [x]° are the constant terms for each reactant from the definition of thermodynamic activity, 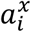:

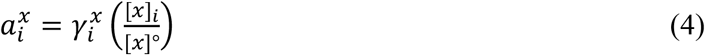

At equilibrium, Δ*G*^rxn^ = 0, and Equation 1 may be reduced to the following relationship:

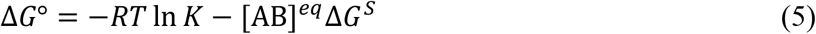

With regard to Equation 5, the equilibrium quotient, *K*, is defined as the ratio of complex to free reactants at equilibrium in the direction of association. As with the classical approach, all concentrations are treated as dimensionless quantities due to division by a reference concentration in the derivation, but the concentrations must be inserted in the same units, typically molarity for dilute solutions.

The derivation of Equation 5 has been questioned by those who contend that the chemical potential of all water molecules in a solution at equilibrium must be equal, and, therefore, Δ*G*^S^ must be zero. However, this notion is misguided. The change in free energy of the system at equilibrium (reactants plus solvent) is zero, but this requirement does not preclude the existence of subpopulations of water molecules that differ in chemical potential due to neighboring solutes (*i*.*e*., boundary conditions). With respect to the solvent, a state of equilibrium means only that the movement of a water molecule from one position to a location of differing chemical potential must be countered by a second water molecule that moves in the opposite direction. Water molecules should be viewed as existing in a dynamic equilibrium between the hydration spheres of the reacting species and all other locations that comprise the bulk phase.^4^

Another common misconception is that all concentration-dependent changes in equilibria should be attributed to changes in the activity coefficients of the reactants. This explanation presumes – often without evidence – that self-interactions (*i*.*e*., reactant A with reactant A) are responsible for the observed changes in equilibria. But does this explanation seem plausible when the reactant concentrations are in the millimolar range or below? For a small reacting species at a concentration of 10 mM, the ratio of water molecules to reactant molecules is approximately 5500:1, and water–reactant interactions should far outnumber reactant–reactant interactions.^3,4^ In the current work, the ratio of excess water to reactant will be reduced due to the excluded volume of the macromolecules, but the employed concentrations of macromolecule are also reduced, rarely exceeding 1.0 mM. Thus, reactant–reactant interactions remain a rare event compared to reactant–solvent interactions, and the activity coefficient of each reactant is expected to be a constant within the tested concentration range.

The current study attributes concentration-dependent changes in binding equilibria to the concomitant changes in solvation. The governing equation, Equation 5, can be tested by measuring the equilibrium quotient as a function of product concentration. If the experimental data agree with the governing equation, then a plot of -*RT*ln*K* versus [AB], the concentration of complex at equilibrium, should yield a straight line with a slope equal to Δ*G*^S^ and a *y*-intercept equal to Δ*G*°. In the work that follows, Equation 5 is applied for the first time to model biomolecular interactions that involve DNA oligonucleotides and proteins.

## MATERIALS AND METHODS

### Reagents

The DNA oligonucleotides utilized in this work are identified in Scheme 1. All oligonucleotides were purchased from Integrated DNA Technologies. The concentrations of stock DNA solutions were checked prior to each ITC run using a Cary 60 spectrophotometer. The DNA buffer consisted of 100 mM NaCl, 0.10 mM EDTA, and 10 mM sodium phosphate, pH 6.8.

**Scheme 1.** Oligonucleotide names, sequences, and extinction coefficients

|  |  |  |
| --- | --- | --- |
| oligo9: | 5'- CAA ATA AAG – 3' | $\epsilon_{260} = 100,900 \text{ M}^{-1} \cdot \text{cm}^{-1}$ |
| oligo9c: | 5'- CTT TAT TTG – 3' | $\epsilon_{260} = 80,000 \text{ M}^{-1} \cdot \text{cm}^{-1}$ |
| oligo10: | 5'- ATG CTG ATG C – 3' | $\epsilon_{260} = 94,800 \text{ M}^{-1} \cdot \text{cm}^{-1}$ |
| oligo10c: | 5'- GCA TCA GCA T – 3' | $\epsilon_{260} = 97,000 \text{ M}^{-1} \cdot \text{cm}^{-1}$ |

Ribonuclease A from bovine pancreas (Fisher, BP2539) and α-lactalbumin from bovine milk (Sigma, L6010) were used for protein studies. Protein concentrations were measured by spectrophotometry using extinction coefficients of 9800 M^-1^·cm^-1^ at 278 nm (RNase A) and 28,500 M^-1^·cm^-1^ at 282 nm (α-lactalbumin). The ribonuclease inhibitor, uridine 3’-monophosphate (3’-UMP), was purchased as a disodium salt from Carbosynth (NU07383), and concentrations were determined by using an extinction coefficient of 9780 M^-1^·cm^-1^ at 262 nm, as reported for 5’-UMP.^5^

### Calorimetry

Isothermal titration calorimetry (ITC) was performed with a Microcal instrument, model VP-ITC, using the analysis programs provided by the manufacturer. All solutions were degassed briefly under vacuum (ThermoVac). Prior to each calorimetry run, the sample cell (∼1.45 mL volume) was cleaned and rinsed with one volume of the corresponding sample buffer. For DNA trials, the injection syringe (∼0.28 mL) was filled with the complementary oligonucleotide at a concentration 20-fold higher than the cell for oligo9 experiments and 10-fold higher for oligo10 experiments. For RNase A trials, stock protein solutions were made fresh and 3’-UMP stocks were stored at -20°C, both in a buffer solution of 25 mM Bis-Tris with 25 mM KCl adjusted to pH 6.0. The sample cell was loaded with the desired RNase A concentration, and the injection syringe contained 3’-UMP at a concentration 10-fold or 20-fold higher than RNase A. The typical ITC protocol consists of 28 injections of 10 μL volume or 55 injections of 5 μL volume. In all trials, the first injection was set at 2 μL, and the first titration peak was discarded from the analysis due to the unavoidable error in the injection volume after filling and moving the syringe to the calorimeter base. A background ITC file, or a constant corresponding to the background injection enthalpy, was subtracted from each integrated peak prior to analysis.

### Circular Dichroism Spectroscopy

A 100 mg/mL stock solution of α-lactalbumin was prepared in 10 mM Tris buffer (pH 7.0) containing 10 mM EDTA and 1.25 M or 3.0 M guanidine HCl (Fisher). The stock protein solution was diluted to the desired concentrations in the same buffer and transferred to cuvettes of varying pathlengths such that the product of c·*l* from Beer’s law was constant. Wavelength scans were carried out on an Aviv Model 215 circular dichroism spectrometer at 25 °C in the near UV region (250-320 nm). Five scans per sample were taken and averaged. A control buffer solution was subtracted from each result before exporting the data to an Excel spreadsheet for analysis.

### Modelling the Governing Equation

Equation 5 was modelled by setting the total concentrations of the ssDNA (X_t_) and its complement (M_t_) as known inputs after each titration point and by using a mass balance expression to replace the concentration of the free ssDNA in the governing equation, where Z is the unknown concentration of duplex at equilibrium:

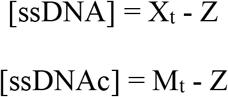

The value of X_t_ after each injection was adjusted by a constant increment, set up as an array of 28 injections of 10 μL volume at a concentration 10-fold higher than the starting concentration of ssDNA oligonucleotide in the cell, M_t_. The two free energy parameters, ΔG° and ΔG^S^, were set as constants that approximate the experimentally-obtained values for oligo10/10c binding. The concentration of bound complex (Z) was solved by WolframAlpha after each injection from the following expression, in accord with Equation 5:

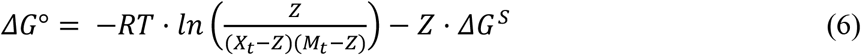

Typically, two or three roots are obtained from WolframAlpha, but only one root is a real number below the maximum possible concentration. After solving for Z, the equilibrium quotient within the natural logarithm term is calculated, as well as the heat released after each injection. The heat may be estimated by multiplying a constant molar binding enthalpy by the increase in Z relative to the previous injection. See Data Availability statement for link to further modelling details.

## RESULTS AND DISCUSSION

In order to estimate the value of Δ*G*^S^ from experimental data, the last term in the Equation 5, [AB]·Δ*G*^S^, must be of sufficient magnitude to observe a shift in the equilibrium. For DNA and proteins, this presents a challenge because, depending on the value of Δ*G*^S^, it may not be possible to push the concentration of the macromolecule high enough to observe an effect. In previous work employing small molecules, the ideal upper concentration varied from 3-10 mM.^3^ With this issue in mind, the following studies were carried out to assess the feasibility of detecting a concentration-dependent change in each equilibrium, as predicted by the governing equation.

### Framework Applied to DNA Hybridization

In the case of DNA hybridization, Equation 5 may be expressed as follows:

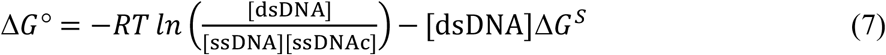

where [dsDNA] is the concentration of double-stranded product at equilibrium, and where [ssDNA] and [ssDNAc] represent the equilibrium concentrations of the single-stranded DNA and its complement, respectively. Because ITC is best reserved for binding affinities in the range of *K* = 10^4^ to 10^7^, short DNA oligonucleotides were employed for the hybridization studies. Figure 1 summarizes the result for the shorter of two model reactions, oligo9 with oligo9c. The measured value of *K* declined modestly from 6.1 x 10^4^ at 0.010 mM duplex to 5.2 x 10^4^ at 0.150 mM duplex. This same duplex has been studied previously, but the prior investigations used a high salt concentration of 1.0 M NaCl which precludes a meaningful comparison of *K* values to the current work in a 10-fold lower salt concentration.^6,7^ As seen in Figure 1, the data for oligo9/9c binding yield a reasonable fit to Equation 7, leading to free energy values of Δ*G*° = -6.52 ± 0.04 kcal/mol and Δ*G*^S^ = +710 ± 500 kcal/mol. The large uncertainty in Δ*G*^S^ is an unavoidable outcome of the narrow concentration range employed.

**Figure 1.**
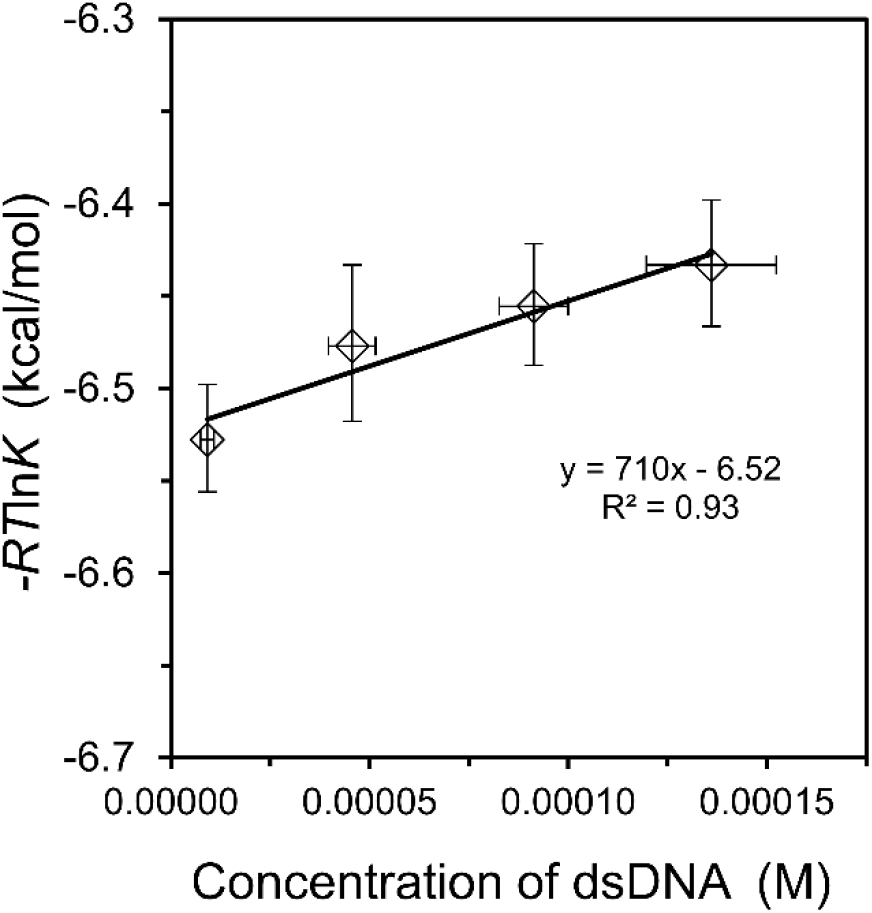
Hybridization of oligo9 with oligo9c at 25 °C. Equilibrium binding quotients were obtained by ITC as a function of concentration. The linear fit is indicated on the graph, from which the slope corresponds to Δ*G*^S^ and the *y*-intercept corresponds to Δ*G*°, in accord with Equation 7. Error bars are based on three or more trials at each concentration. See Table S1 for corresponding values of *K* and Δ*H*^ITC^.

In a second test of hybridization, a DNA duplex of stronger binding affinity was employed. Oligo10 and oligo10c form five G:C base pairs, as opposed to just two for oligo9/9c, and a binding affinity on the order of 10^6^ has been reported for this duplex, as measured by ITC under similar conditions.^8,9^ In the current study, the concentration range was expanded to an upper value of 0.200 mM, and the ITC experiments were performed at four different temperatures. As shown in Figure 2, a reasonable fit to Equation 7 was observed at each temperature. For the specific dataset at 25 °C, the value of *K* decreased from 7.72 x 10^6^ at the lowest concentration to 1.79 x 10^6^ at the highest concentration, a much broader range than observed with oligo9/9c.

**Figure 2.**
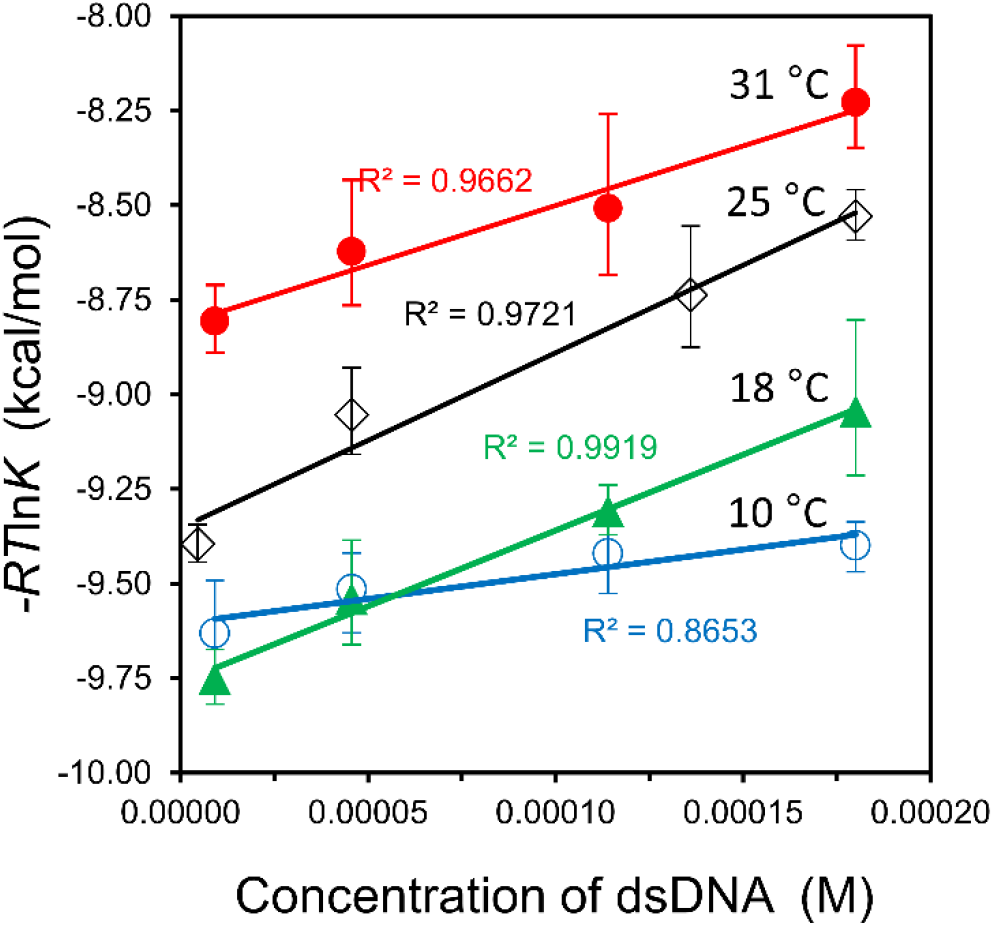
Hybridization of oligo10 with oligo10c. The corresponding temperatures and goodness of fits to Equation 7 are shown on the graph. Note that the starting conformations of the oligonucleotides vary with temperature due to different amounts of intramolecular stacking. See Table S2 for corresponding values of *K* and Δ*H*^ITC^.

These *K* values bookend a reported value of 3.5 x 10^6^, as obtained from the average of multiple measurements in the concentration range of 0.004-0.051 mM.^9^

The intersection of datasets at 10 °C and 18 °C in Figure 2 was initially viewed as curious and unexpected. After further reflection, however, it was realized that the datasets in Figure 2 are influenced by intramolecular base stacking within each oligonucleotide which also varies with temperature (Figure 3a). This residual structure can be detected and quantified by differential scanning calorimetry.^9–11^ Thus, the starting point for hybridization is different at each temperature; a lower temperature begins with more intramolecular stacking than a higher temperature.

**Figure 3.**
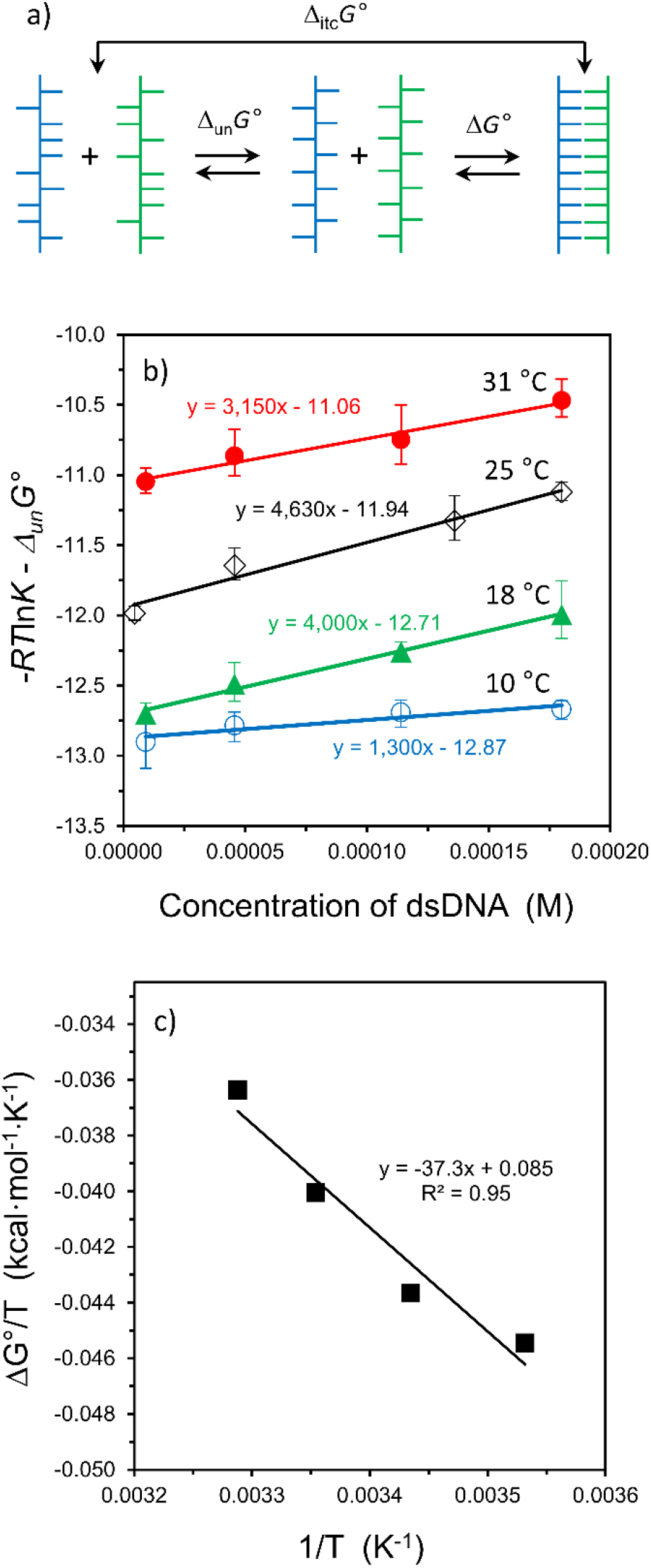
Revised analysis of oligo10/10c hybridization. (a) Schematic diagram of two-step process for unstacking of intramolecular bases prior to duplex formation; Δ_itc_*G*° = Δ_un_*G*° + Δ*G*°. (b) Updated graph for hybridization of oligo10/10c using Δ*G*° values in Table 1. (c) A van’t Hoff analysis of the Δ*G*° values in Table 1.

To place the temperature-dependent results on the same starting basis, the free energy required to unstack each oligonucleotide (Δ_un_*G*°) was calculated using the reported standard state values for the enthalpy and entropy of unstacking for oligo10 and oligo10c.^9,12^ The values of Δ*G*° obtained after subtracting the free energy of unstacking from the ITC-measured value (Δ_itc_*G°*) are summarized in Table 1.

**Table 1.** Accounting for the free energy of unstacking.

| $T$<br>(°C) | $\Delta_{\text{un}}G^\circ = \Delta_{\text{un}}H^\circ - T\Delta_{\text{un}}S^\circ$ | | | Fraction<br>Stacked ($f$) | $\Delta_{\text{itc}}G^\circ$<br>from<br>Fig. 2 | $\Delta G^\circ =$<br>$\Delta_{\text{itc}}G^\circ - f\Delta_{\text{un}}G^\circ$ |
| --- | --- | --- | --- | --- | --- | --- |
|  | oligo10 | oligo10c | Sum |  |  |  |
| 10 | +1.62 | +1.65 | +3.27 | 1.0 | -9.60 | -12.87 |
| 18 | +1.45 | +1.50 | +2.95 | 1.0 | -9.76 | -12.71 |
| 25 | +1.30 | +1.37 | +2.67 | 0.97 | -9.35 | -11.94 |
| 31 | +1.18 | +1.26 | +2.44 | 0.92 | -8.82 | -11.06 |

1. Free energy units in kcal/mol. 2. Values of $\Delta_{\text{un}}H^\circ$ and $\Delta_{\text{un}}S^\circ$ from Lang and Schwarz.<sup>9</sup> 3. Fraction in stacked conformation estimated from Zhou, et al.<sup>12</sup>

Presumably, the temperature-specific value of Δ*G*^S^ is also influenced by the unstacking transition of ssDNA, but the concentration dependence of the unstacking transition is unknown for oligo10 and oligo10c and, due to an expected weak signal, would be difficult to measure by differential scanning calorimetry at the lower concentrations utilized in this study. For this reason, the datasets in Figure 2 were re-plotted in Figure 3(b) to reflect the new *y*-intercepts (Δ*G*°) without altering the slopes (Δ*G*^S^). The datasets at 10 °C and 18 °C no longer intersect in the revised graph, and a van’t Hoff analysis of the Δ*G*° values leads to a hybridization enthalpy of Δ*H*° = -37 kcal/mol and entropy of Δ*S*° = -0.085 kcal·mol^-1^·K^-1^, as shown in Figure 3(c). These energies correspond to oligonucleotides that start in a fully-unstacked conformation. The Δ*H*° value is smaller than the ITC-measured enthalpy of -47 ± 5 kcal/mol, as obtained from the average of all trials at 25 °C.

Note that the slopes of the lines in Figure 3(b) do not follow a meaningful trend in magnitude. This is likely due to the lack of a correction for intramolecular stacking as a function of concentration for Δ*G*^S^. In general, all hybridization experiments yield a large and positive slope, indicating an unfavorable change in solvation free energy upon duplex formation. For oligo10/10c binding at 25 °C, Δ*G*^S^ = +4,600 kcal/mol (+460 kcal/mol per base pair).Volumetric measurements for a duplex of similar size and composition to oligo10/10c led to an estimate of ∼180 water molecules that are released upon hybridization.^13^ Thus, many water molecules may contribute to the sign and magnitude of Δ*G*^S^ for this model hybridization reaction. Furthermore, duplex formation may be a case where the water that remains bound to the reaction product also contributes significantly to the change in solvation free energy. Formation of the double-helix is accompanied by a “spine” of hydration in the minor groove, as detected in solution by NMR at 4-10 °C^14^ and by nonlinear vibrational spectroscopy at room temperature.^15^ The degree of hydrogen bonding between water molecules along the spine is known to be weaker for AT-rich tracts relative to GC-rich tracts in dsDNA,^15^ providing one possible reason for the differing values of Δ*G*^S^ per base pair when comparing oligo9/9c to oligo10/10c. In both cases, the unfavorable change in solvation free energy implicates the displacement of water molecules as a major driving force for interaction of DNA-binding drugs and proteins.^16,17^

### Modelling of Titration Experiments

One nuance of the thermodynamic framework employed here is that it predicts the equilibrium quotient should change after each injection during the progress of an isothermal titration experiment.^3^ To demonstrate this phenomenon, a modelling study was performed with experimental parameters that approximate the hybridization of oligo10/10c at 0.20 mM concentration (Figure 4). When the change in solvation free energy is positive, the equilibrium quotient declines after each injection for the first half of the titration, approaching a constant value near the midpoint of the titration, as shown in Figure 4(a). The midpoint corresponds to a 1:1 molar ratio in the total concentration of each oligonucleotide. The last half of the titration is characterized by a nearly-constant value of *K* because the oligonucleotide that started in the calorimeter cell has become the limiting reactant and because nearly all of this limiting reactant has formed a duplex. Simulated ITC enthalpy curves for Δ*G*^S^ values of zero and +4000 kcal/mol are shown in Figure 4(b). Note how the curve for Δ*G*^S^ = +4000 (red solid line) coincides with a curve obtained from the classical equation when *K* is a constant (gray dots). This constant *K* value corresponds to the last injection prior to exceeding 1:1 molar ratio in the simulated curve for Δ*G*^S^ = +4000 in Figure 4(a). For this reason, the *K* values obtained from the ITC instrument’s fitting program, which utilizes the classical equation for binding equilibria, are matched with the concentration of the limiting reactant in the calorimeter cell near the 1:1 point, as detailed in a previous study.^3^ An experimental ITC run at the same concentration as the simulation may be found in Figure S1(d).

**Figure 4.**
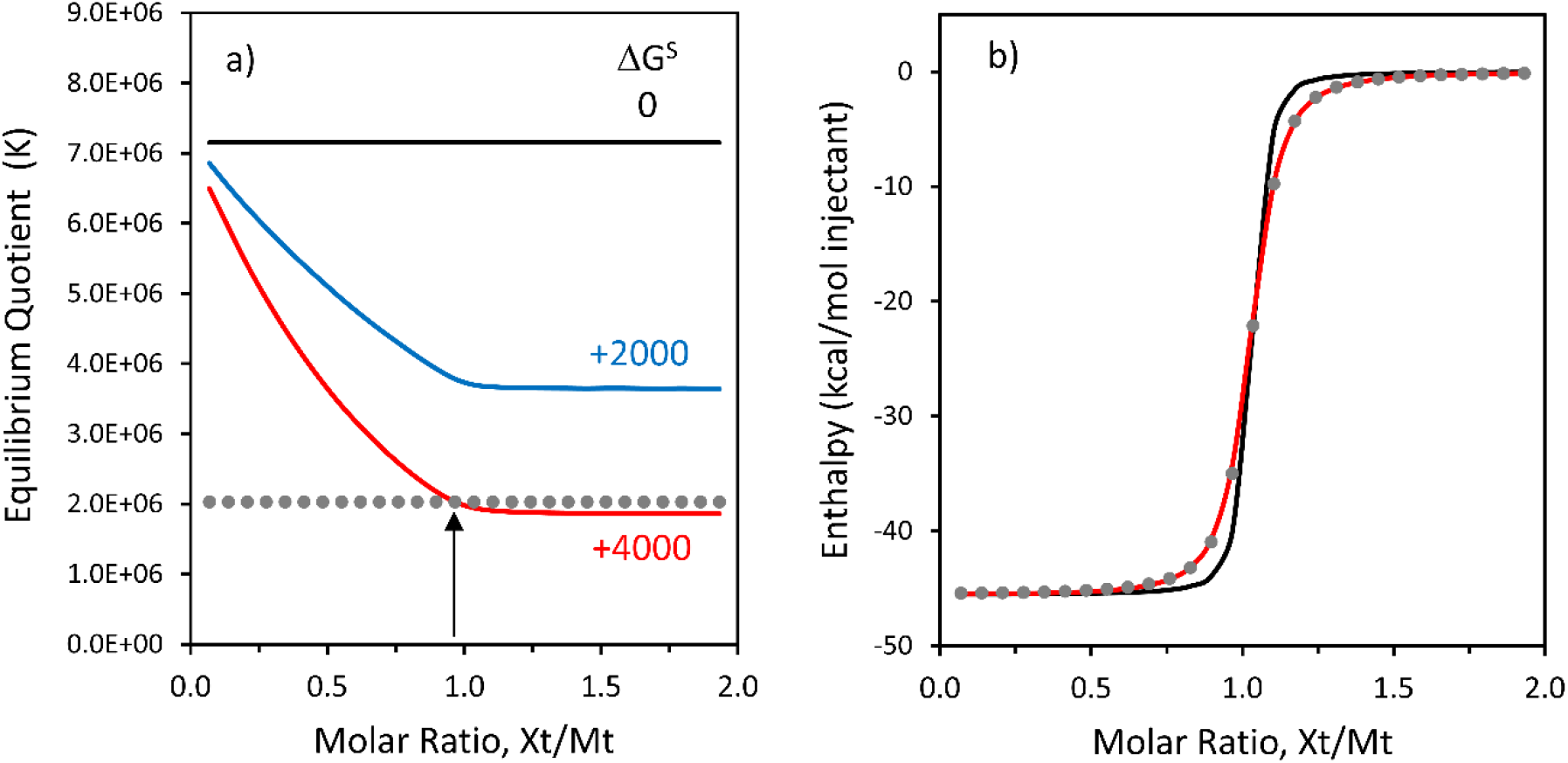
Modelling of titration results for oligo10/10c at 0.20 mM concentration and 25 °C. (a) Equilibrium quotients as a function of Δ*G*° and Δ*G*^S^, as obtained from Equation 6 using WolframAlpha to calculate the unknown concentration of duplex after each injection. The *x*-axis represents the molar ratio of total injected species (X_t_) to total binding partner in the ITC cell (M_t_). For the three solid curves, Δ*G*° = -9.30 kcal/mol, and Δ*G*^S^ is given next to each curve in kcal/mol. Simulations for Δ*G*^S^ = 0 correspond to a constant value of *K*. The gray dotted line represents a constant *K* value of 2.03 x 10^6^, and the vertical arrow indicates the intersection with the dataset for Δ*G*^S^ = +4000, just prior to exceeding a molar ratio of 1.0. (b) Simulated ITC enthalpy curves corresponding to datasets in panel (a). The gray dots represent *K* = 2.03 x 10^6^ (Δ*G*^S^ = 0, Δ*G*° = -8.60 kcal/mol) and nearly coincide with the red curve (for which Δ*G*^S^ = +4000, Δ*G*° = -9.30 kcal/mol).

### Framework Applied to Protein–Ligand Interactions and Protein Folding

Ribonuclease A (RNaseA) has served as a model protein for enzyme structure, enzyme function, and inhibitor binding studies for decades.^18,19^ A classical ITC study of protein–ligand interactions, carried out by J.F. Brandts and coworkers, examined the binding of RNaseA with the nucleotide inhibitor 2’-CMP and noted a significant decrease in the equilibrium ratio with increasing reactant concentration.^20^ This rare example of a concentration-dependent equilibrium was reported, in part, because the authors wanted to demonstrate the versatility of a new ITC instrument over a 50-fold range in protein concentration. The authors hypothesized that the concentration-dependent *K* values were due to dimerization or aggregation of RNaseA, though a footnote in a subsequent report attributed the results to 2’-CMP “if they arise from nonideality effects.”^21^

In another study employing RNaseA, a salt-dependent change in binding affinity with the inhibitor 3’-UMP was observed, but the investigators increased the protein concentration simultaneously, making it impossible to separate solvation effects from salt effects.^22^ In the current work, binding of 3’-UMP to RNaseA is re-examined at five concentrations while maintaining a constant NaCl concentration (Figure 5, upper line). For comparison, the 2’-CMP binding result from the Brandts study is plotted on the same graph (Figure 5, lower line). Because some of the experimental details are unknown for the Brandts dataset, it was assumed that the concentration of complex at the 1:1 molar titration point (*x*-value on graph) is 90% of the starting concentration of RNaseA in the calorimeter cell.

**Figure 5.**
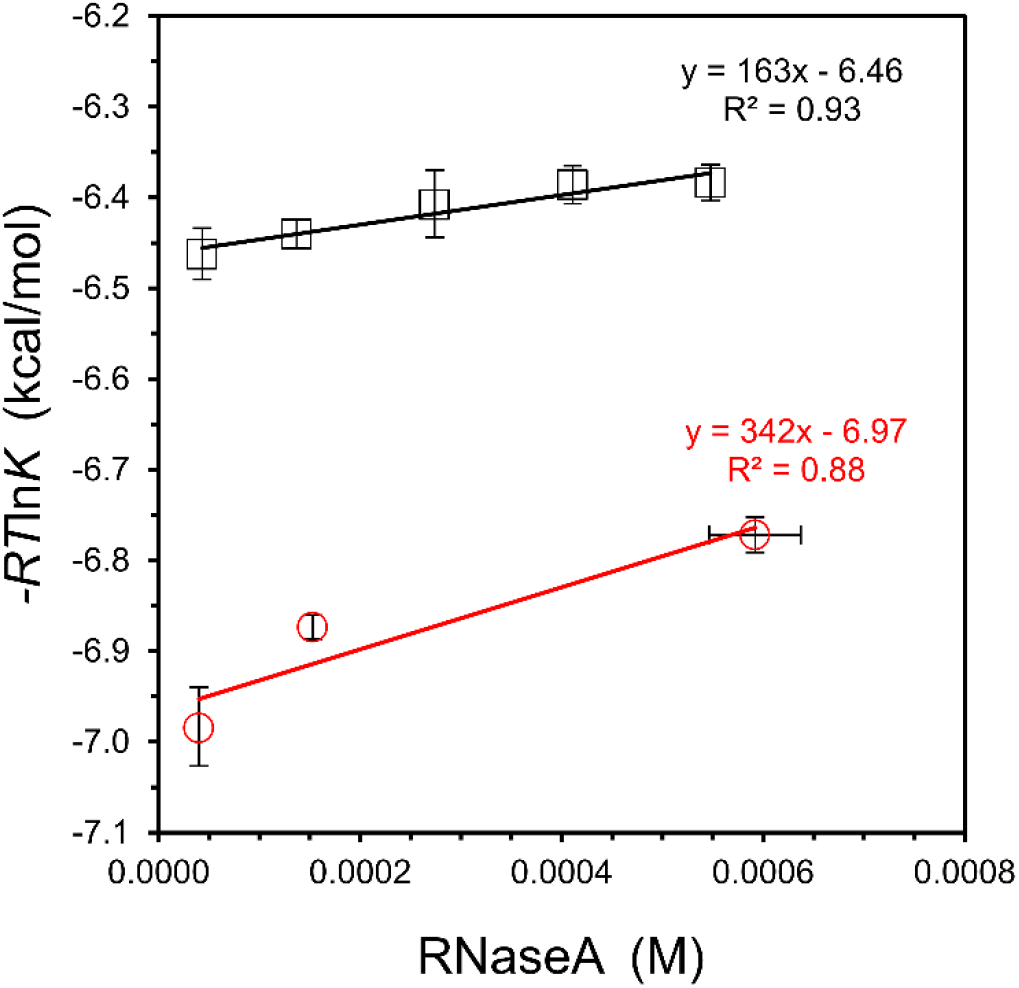
Binding analysis of 3’-UMP/RNaseA and 2’-CMP/RNaseA as a function of concentration. Binding of the 3’-UMP inhibitor was measured in 25 mM KCl and 25 mM BisTris buffer, pH 6.0, at 25 °C (top, black squares). The *K* values for 2’-CMP are tabulated in Wiseman, et al., as measured in 200 mM KCl and 200 mM potassium acetate buffer, pH 5.5, at 28 °C (bottom, red circles).^20^ Error bars are based on four or more trials for both datasets. For 3’-UMP, see Table S3 for corresponding values of *K* and Δ*H*^ITC^.

Although a concentration-dependent trend is difficult to establish with high certainty in such a narrow range of protein concentration (highest concentration of 0.6 mM), a linear fit is observed in Figure 5 for 3’-UMP/RNaseA, leading to binding free energies of Δ*G*^S^ = +160 ± 100 kcal/mol and Δ*G*° = -6.46 ± 0.03 kcal/mol. The estimated change in solvation energy for 2’-CMP binding is about twice that of 3’-UMP, but the number and distribution of tested concentrations are not ideal, leading to a weaker fit to the governing equation for the 2’-CMP study (Figure 5, bottom line). For both nucleotide inhibitors, the change in equilibrium with reactant concentration is viewed as the outcome of accounting for the change in solvation energy, in accord with Equation 5. An unfavorable change in solvation energy for nucleotide/RNaseA binding seems consistent with (1) a volumetric study that estimates ∼210 water molecules are released to the bulk upon binding^23^ and (2) crystal structures that indicate up to 10 water molecules (of low entropy) remain hydrogen-bonded to the inhibitor in the complex.^24,25^

In the last example, a conformational equilibrium is tested against the governing equation. Protein folding is often approximated as a 2-state process, U ↔ F, where U is the unfolded or denatured state and F is the folded, native state of the protein. The 2-state assumption holds well for many smaller proteins, and reconfiguration of Equation 5 to reflect a 2-state folding equilibrium leads to the following expression:

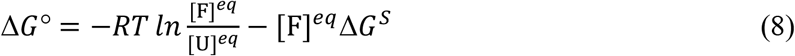

One challenge in applying Eq. 8 to protein folding equilibria is the strong cooperativity of most folding transitions. In order to detect a concentration-dependent change in the equilibrium of a model protein, the structure of α-lactalbumin (123 amino acid residues) was examined in 1.25 M guanidinium chloride (GuHCl) by using circular dichroism in the near-UV region to follow changes in tertiary structure. In this case, the unfolded state is a molten globule that retains some of its secondary structure. A concentration of 1.25 M GuHCl is near the midpoint of a 2-state transition for the Ca^2+^-free apoprotein when following the CD spectrum at 270 nm.^26^ To ensure that all protein is in the apo-state, it was critical to include a higher-than-normal EDTA concentration of 10 mM in the stock solution (∼7 mM protein) to chelate residual calcium ion from the freeze-dried protein preparation.

As seen in Figure 6 below, a concentration-dependent change in tertiary structure is not detected in 1.25 M GuHCl; the changes in CD spectra are negligible when varying the protein concentration from 1-100 mg/mL. This observation implies that the equilibrium between the folded and unfolded state does not change significantly with concentration. A similar, concentration-independent spectrum is observed for α-lactalbumin in 3 M GuHCl (Figure S3). Of possible relevance, it has been suggested that Δ*G*^S^ ≈ 0 for hydrophobic interactions in general, as exemplified by the binding of small nonpolar molecules to β-cyclodextrin.^3,4^ Thus, the concentration-independent result observed for the conformational equilibrium of α-lactalbumin is consistent with the concept that the folded state of a protein is stabilized primarily by nonpolar interactions and the hydrophobic effect, as often ascribed.^27^

**Figure 6.**
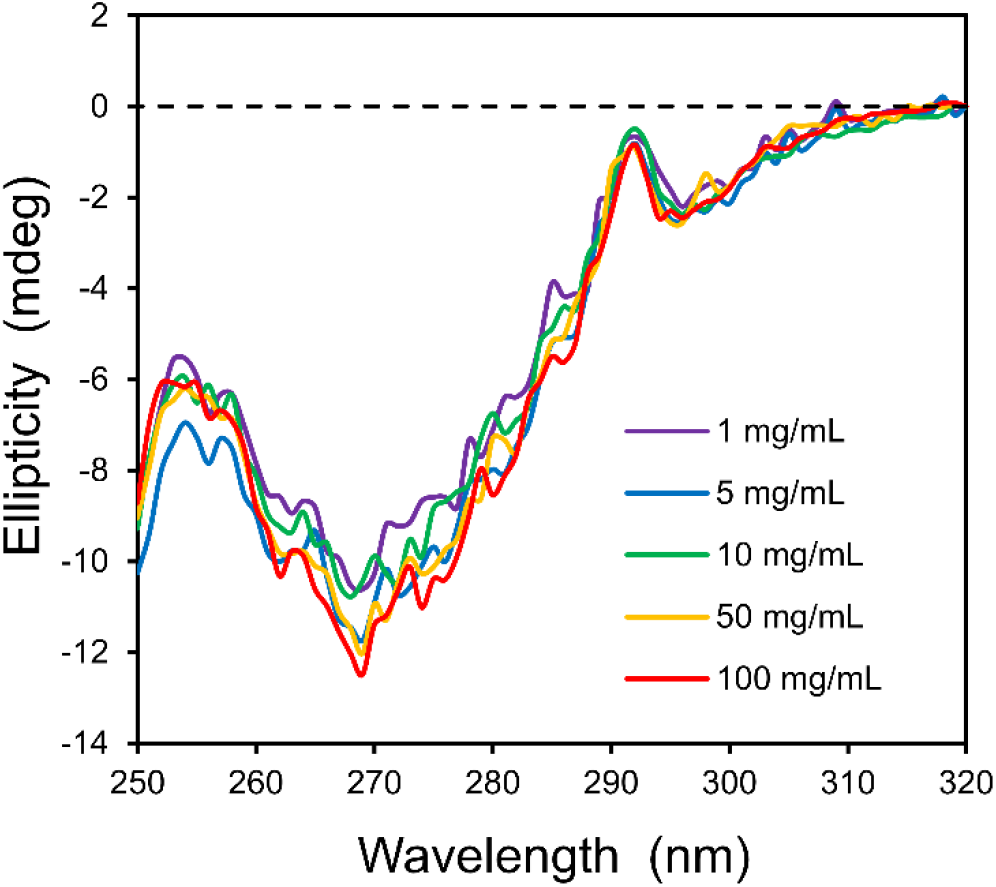
The two-state equilibrium for α-lactalbumin in guanidinium chloride does not change with protein concentration. The near-UV CD profiles of α-lactalbumin in 1.25 M GuHCl, 10 mM EDTA, and 10 mM Tris at 25 °C are shown as a function of protein concentration. Note that the path length of the cuvet (*l*) was varied from 0.01 – 1.0 cm to maintain a constant number of protein molecules in the path of the light source (c·*l* = 1 cm·mg/mL).

## CONCLUSIONS

The experiments in this investigation demonstrate the feasibility of measuring concentration-dependent changes in equilibria for DNA hybridization and protein–ligand binding. In the case of DNA hybridization, a shift in each tested equilibrium was detectable because the change in solvation energy is large enough in magnitude to overcome the low, sub-millimolar concentrations of DNA employed. A value for Δ*G*^S^ of +460 kcal/mol/base pair, as obtained for oligo10/10c binding, indicates that the solvation shell of dsDNA is thermodynamically unfavorable relative to the bulk phase and implicates the displacement of water molecules as a major energetic contributor for DNA-binding proteins and drugs.

The interaction of ribonuclease A with 3’-UMP was tested as a model for protein–ligand binding and compared to a past result from the literature that used a different nucleotide inhibitor. Binding of both inhibitors led to a positive, unfavorable change in solvation energy when analyzed by Equation 5. Detecting a change in equilibrium was more challenging for this model system because of the practical limits in maximum protein concentration, coupled with the relatively low value of Δ*G*^S^ (+160 kcal/mol for 3’-UMP). A conformational study with α-lactalbumin did not detect a change in equilibrium with increasing concentration, but this result is expected if a near-zero change in solvation energy is a general characteristic of hydrophobic interactions and if hydrophobic interactions are the dominant force in protein folding equilibria.

This work suggests a fundamental change in the application of thermodynamics to biological reaction equilibria. The scientific community is encouraged to examine other macromolecular interactions as a function of concentration to test further the utility of the governing equations employed here. Computational methods that ascertain the chemical potential of water molecules as a function of position,^28^ both adjacent to and remote from a solute boundary, may supplement this experimental approach and facilitate a deeper appreciation for the role of solvation energy in molecular recognition.

## ASSOCIATED CONTENT

### Supporting Information

The following files are available free of charge: table of thermodynamic values for each model binding pair, sample ITC curves, CD spectra of α-lactabumin in 3 M GuHCl.

## AUTHOR INFORMATION

