## Supporting Information for "Solvation Free Energy in Governing Equations for DNA Hybridization, Protein–Ligand Binding, and Protein Folding"

**Page**

Table S1 Thermodynamic Values and Uncertainties for Oligo9/9c ........................................ S2

Table S2 Thermodynamic Values and Uncertainties for Oligo10/10c ..................................... S3

Table S3 Thermodynamic Values and Uncertainties for 3'-UMP Binding with RNase A ......... S4

Figure S1 Sample ITC Curves for Oligo10/10c Duplex Formation ........................................... S5

Figure S2 Sample ITC Curves for 3'-UMP/RNaseA Binding ..................................................... S6

Figure S3 Near-UV CD Profiles of $\alpha$-Lactalbumin in 3.00 M GdHCl ......................................... S7

**Table S1.** Thermodynamic Values and Uncertainties for Oligo9/9c

| **Oligo9/9C** | | **25 °C** | |
|---|---|---|---|
| Cell Conc (mM) | | *K* ± Error ($\times 10^4$) | $\Delta H^{ITC}$ kcal/mol |
| Start | at 1:1 | | |
| 0.0100 | 0.0091 | 6.07 ± 0.33 | -45 |
| 0.0500 | 0.0456 | 5.57 ± 0.36 | -48 |
| 0.100 | 0.0913 | 5.36 ± 0.29 | -49 |
| 0.150 | 0.136 | 5.17 ± 0.27 | -48 |
| Free Energies: (kcal/mol) | | $\Delta_{itc}G°$ = -6.52 ± 0.04 | |
| | | $\Delta G^S$ = +710 ± 500 | |

**Table S2.** Thermodynamic Values and Uncertainties for Oligo10/10c

| **Oligo10/10C** | | **10 °C** | | **18 °C** | |
|---|---|---|---|---|---|
| Cell Conc (mM) | | *K* ± Error (x $10^6$) | $\Delta H^{ITC}$ kcal/mol | *K* ± Error (x $10^6$) | $\Delta H^{ITC}$ kcal/mol |
| Start | at 1:1 | | | | |
| 0.0100 | 0.00913 | 27.2 ± 7.7 | -40 | 20.9 ± 2.6 | -42 |
| 0.0500 | 0.0456 | 22.1 ± 4.1 | -41 | 14.5 ± 3.4 | -40 |
| 0.125 | 0.114 | 18.7 ± 3.2 | -42 | 9.72 ± 1.11 | -37 |
| 0.200 | 0.180 | 18.0 ± 2.1 | -38 | 6.16 ± 2.08 | -38 |
| Free Energies: (kcal/mol) | | $\Delta_{itc}G^\circ$ = -9.60 ± 0.18 | | $\Delta_{itc}G^\circ$ = -9.76 ± 0.14 | |
| | | $\Delta_{itc}G^S$ = +1300 ± 1200 | | $\Delta_{itc}G^S$ = +4000 ± 1600 | |

| **Oligo10/10C** | | **25 °C** | | **31 °C** | |
|---|---|---|---|---|---|
| Cell Conc (mM) | | *K* ± Error (x $10^6$) | $\Delta H^{ITC}$ kcal/mol | *K* ± Error (x $10^6$) | $\Delta H^{ITC}$ kcal/mol |
| Start | at 1:1 | | | | |
| 0.0050 | 0.00456 | 7.72 ± 0.64 | -49 | — | — |
| 0.0100 | 0.00913 | — | — | 2.13 ± 0.31 | -49 |
| 0.0500 | 0.0456 | 4.34 ± 0.83 | -48 | 1.57 ± 0.42 | -46 |
| 0.125 | 0.114 | — | — | 1.30 ± 0.44 | -51 |
| 0.150 | 0.136 | 2.54 ± 0.67 | -45 | — | — |
| 0.200 | 0.180 | 1.79 ± 0.20 | -44 | 0.818 ± 0.176 | -44 |
| Free Energies: (kcal/mol) | | $\Delta_{itc}G^\circ$ = -9.35 ± 0.07 | | $\Delta_{itc}G^\circ$ = -8.82 ± 0.10 | |
| | | $\Delta_{itc}G^S$ = +4630 ± 650 | | $\Delta_{itc}G^S$ = +3150 ± 1300 | |

**Table S3.** Thermodynamic Values and Uncertainties for 3'-UMP Binding to Ribonuclease A

| **RNaseA/UMP** | | **25 °C** | |
|---|---|---|---|
| Cell Conc (mM) | | *K* ± Error (x $10^4$) | $\Delta H^{ITC}$ kcal/mol |
| Start | at 1:1 | | |
| 0.0450 | 0.0431 | 5.46 ± 0.26 | -12.6 |
| 0.150 | 0.137 | 5.26 ± 0.14 | -12.6 |
| 0.300 | 0.274 | 4.98 ± 0.31 | -12.2 |
| 0.450 | 0.411 | 4.80 ± 0.17 | -12.2 |
| 0.600 | 0.548 | 4.78 ± 0.16 | -12.4 |
| Free Energies: (kcal/mol) | | $\Delta G°$ = -6.46 ± 0.03 | |
| | | $\Delta G^S$ = +160 ± 100 | |

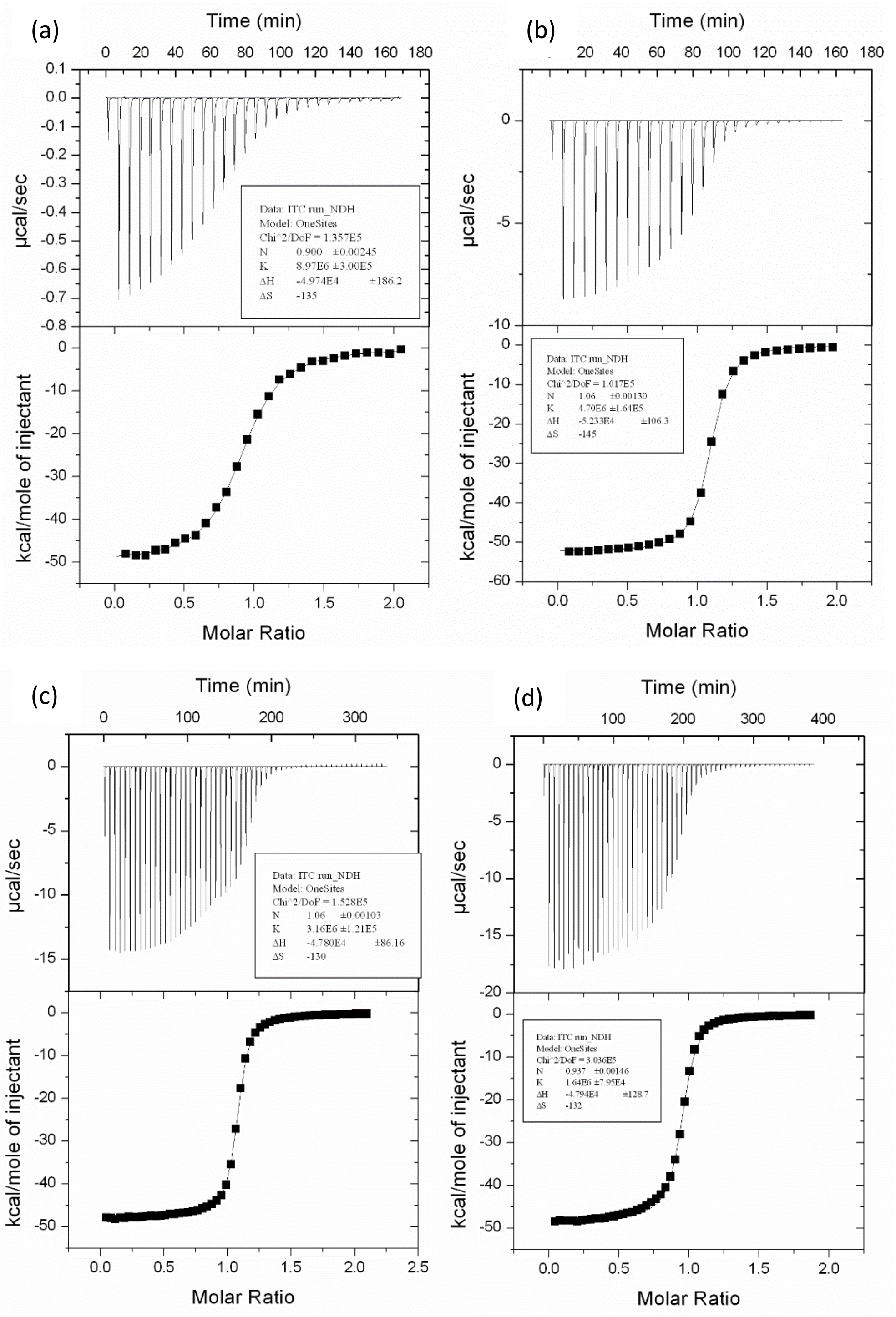


**Figure S1.** Sample ITC curves for oligo10/10c duplex formation. Trials completed at 25 °C with following concentrations of oligo10 in calorimeter cell: (a) 0.005 mM; (b) 0.050 mM; (c) 0.15 mM; and (d) 0.20 mM. The curves in (a) and (b) employ 28 injections of 10 μL, whereas the curves in (c) and (d) employ 55 injections of 5 μL.

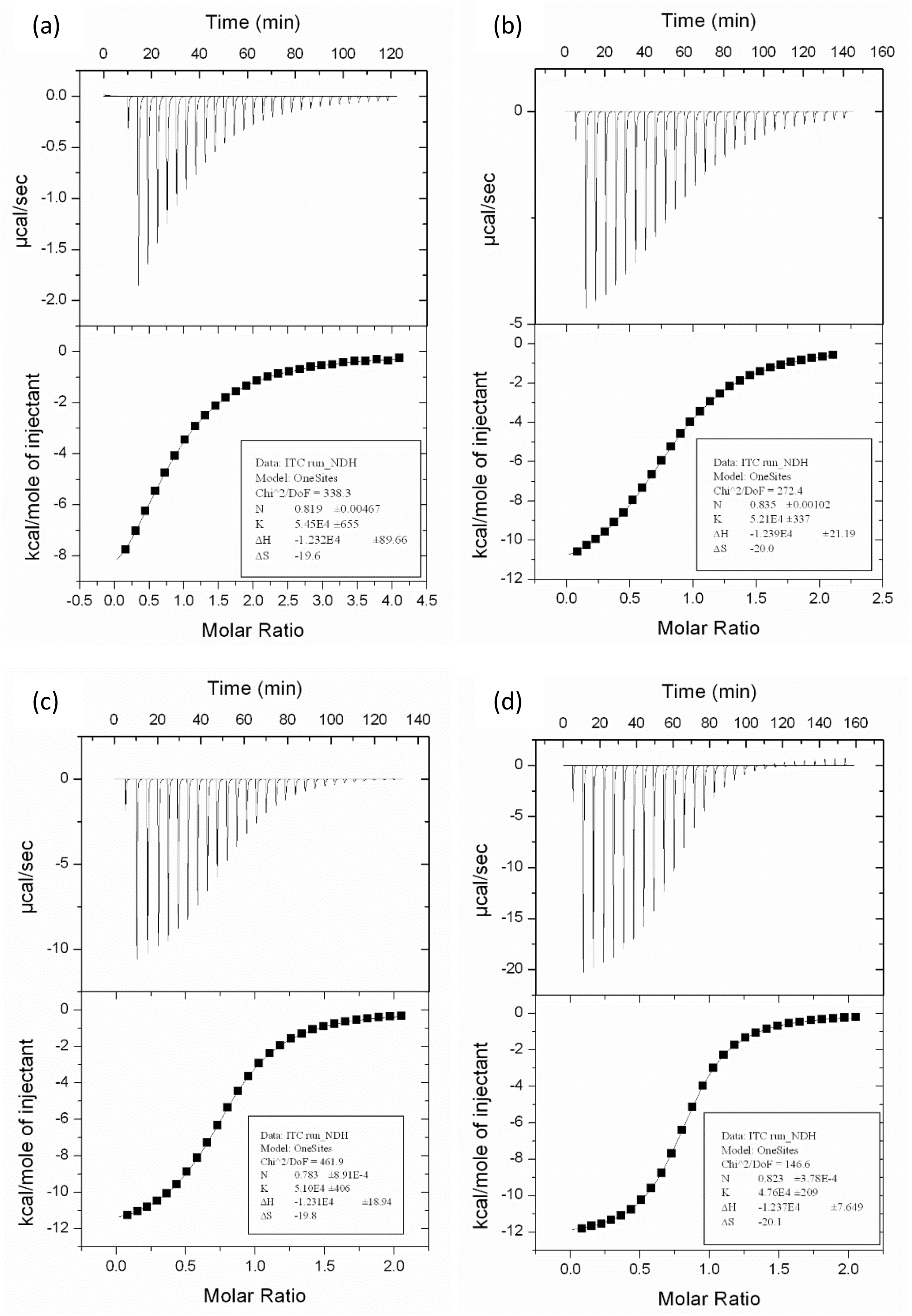


**Figure S2.** Sample ITC curves for 3'-UMP/RNaseA binding. Trials completed at 25 °C with following concentrations of oligo10 in calorimeter cell at start: (a) 0.045 mM; (b) 0.150 mM; (c) 0.300 mM; and (d) 0.600 mM. For 0.045 mM trial only, the 3'-UMP concentration in syringe is 20 x cell concentration.

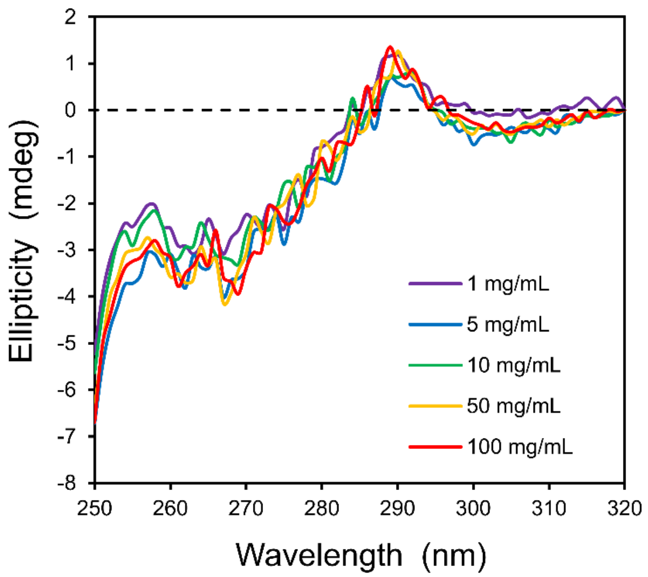


**Figure S3.** Near-UV CD profiles of $\alpha$-lactalbumin in 3.00 M GdHCl. Further data suggesting that the 2-state equilibrium for $\alpha$-lactalbumin in guanidinium chloride does not change with protein concentration. Spectra obtained in 3.00 M GdHCl, 10 mM EDTA, and 10 mM Tris at 25 °C at protein concentrations noted on graph. As done for Fig. 6 in the main text, the path length of the cuvet (*l*) was varied from 0.01 – 1 cm to maintain a constant number of protein molecules in the pathe of the light source ($c \cdot l = 1$ cm·mg/mL).
